# Neuronal gain modulability is determined by dendritic morphology: a computational optogenetic study

**DOI:** 10.1101/096586

**Authors:** Sarah Jarvis, Konstantin Nikolic, Simon R Schultz

**Affiliations:** Centre for Neurotechnology and Department of Bioengineering, Imperial College London, London, UK; Centre for Bio-Inspired Technology and Department of Electrical & Electronic Engineering, Imperial College London, London, UK

## Abstract

The mechanisms by which the gain of the neuronal input-output function may be modulated have been the subject of much investigation. However, little is known of the role of dendrites in neuronal gain control. New optogenetic experimental paradigms based on spatial profiles or patterns of light stimulation offer the prospect of elucidating many aspects of single cell function, including the role of dendrites in gain control. We thus developed a model to investigate how competing excitatory and inhibitory input within the dendritic arbor alters neuronal gain, incorporating kinetic models of opsins into our modeling to ensure it is experimentally testable. To investigate how different topologies of the neuronal dendritic tree affect the neuron’s input-output characteristics we generate branching geometries which replicate morphological features of most common neurons, but keep the number of branches and overall area of dendrites approximately constant. We found a relationship between a neuron’s gain modulability and its dendritic morphology, with neurons with bipolar dendrites with a moderate degree of branching being most receptive to control of the gain of their input-output relationship. The theory was then tested and confirmed on two examples of realistic neurons: 1) layer V pyramidal cells - confirming their role in neural circuits as a regulator of the gain in the circuit in addition to acting as the primary excitatory neurons, and 2) stellate cells. In addition to providing testable predictions and a novel application of dual-opsins, our model suggests that innervation of all dendritic subdomains is required for full gain modulation, revealing the importance of dendritic targeting in the generation of neuronal gain control and the functions that it subserves. Finally, our study also demonstrates that neurophysiological investigations which use direct current injection into the soma and bypass the dendrites may miss some important neuronal functions, such as gain modulation.

**Author Summary:**

- Gain modulability indicated by dendritic morphology
- Pyramidal cell-like shapes optimally receptive to modulation
- All dendritic subdomains required for gain modulation, partial illumination is insufficient
- Computational optogenetic models improve and refine experimental protocols

## Introduction

Neuronal gain modulation occurs when the sensitivity of a neuron to one input is controlled by a second input. Its role in neuronal computation has been the subject of much investigation [1–4], and its dysfunction has been implicated in a range of disorders from attention deficit disorders, through to schizophrenia, autism and epilepsy [5–8]. Neocortical neurons vary in modulability, with gain modulation having been observed in cortical pyramidal cells from layers 2/3, 5 and 6 [9,10], whereas input-output relationships in some other cell types, such as entorhinal stellate cells, appear to be much less modulable [11]. Despite their role as the principal excitatory neuronal class within the cortex, it is unknown which properties of pyramidal cells are necessary in order to modulate their gain.

Gain modulation is signified by a change in the gradient of the input-output function of a neuron, in comparison to an overall change in excitability, which is instead evident as a lateral shift. There have been several proposed mechanisms for how a neuron alters the relationship between its input and output, including the use of shunting inhibition to shift the input-output curve [12,13], and varying the rate of background synaptic noise, decreasing the ability of the neuron to detect target input signals [14,15]. A subsequent theoretical study posits that both mechanisms may be necessary [16], which is supported by experimental evidence from intracellular *in vivo* recordings [17], indicating that these processes are not mutually exclusive and may instead operate in different regimes. Notably, theoretical studies have used point neuron models, while experimental studies have injected current into the soma. However, as in *situ* the processing of individual synaptic inputs occurs within the dendrites rather than somatically, this raises another possibility: that gain modulation may involve dendritic processing. The modulation of gain is affected by the balance between excitation and inhibition, and as dendrites act to integrate inputs from throughout their arbors, their capacity for mediating between attenuation and saturation is highly dependent upon the local configuration of dendritic segments and synaptic inputs [18–24]. This suggests the possibility that the morphology of the dendritic tree itself is sufficient for managing attenuation and saturation of inputs, thereby facilitating a neuron’s capacity for gain modulation.

To date, technical limitations in observing and manipulating activity at multiple locations throughout the dendritic arbor have made experimental studies of the dendritic contribution to neuronal gain control infeasible. While recording from single or a small number of dendritic locations is possible [25], this technique is not suited to manipulating activity over multiple locations, mimicking the thousands of inputs a pyramidal cell receives in *vivo.* However, optogenetics may prove to be a better method for manipulating dendritic activity, as light-activated opsins can be expressed throughout the entire membrane of the neuron – including the dendrites. The existence of both excitatory [26] and inhibitory opsins [27,28] suggests the possibility of altering the balance of excitatory and inhibitory currents locally in dendrites, to act as a synthetic substitute for the effect of excitatory and inhibitory presynaptic input. This raises the prospect of a viable experimental method with which to investigate the mechanisms of neuronal gain modulation in the whole cell, as opposed to studying somatic effects alone.

Here we demonstrate through a computational model that neuronal gain modulation can be determined by cell morphology, by means of a set of dendritic morphological features which mediate between attenuation and shunting to modulate neuronal output. The local interaction of competing excitatory and inhibitory inputs is sensitive to the placement of dendritic sections. This indicates that gain modulation can be achieved by altering the overall balance of excitation and inhibition that a neuron receives, rather than being dependent on the statistical properties of the synaptic input. As experimental validation of our work would require optogenetics, we tested our hypothesis in detailed, biophysical models of opsin-transfected neurons, using experimentally fitted models of channelrhodopsin-2 (ChR2) and halorhodopsin (NpHR), which when activated produced excitatory and inhibitory photocurrents.

This study proposes a new perspective on the contribution of dendritic morphology to the characteristics of neurons, relating the shape of their dendritic arbors directly to their functional role within the neural circuit. We show that all dendritic subdomains are required for gain modulation to occur, suggesting that distinct innervation through synaptic targeting by discrete presynaptic populations could be an effective mechanism by which a neuron’s output can be quickly gated between high gain and low gain modes. By incorporating kinetic opsin models within our detailed, compartmental neuron modeling approach, we make predictions which are directly testable by optogenetic experiments. In particular, our model leads to the proposal of a new illumination protocol for a more naturalistic method of neuronal photostimulation - in which rather than simply imposing spikes or shutting down neuronal activity entirely, it is possible to increase or decrease the gain of a neuron’s response to existing inputs.

We would also like to clarify the meaning of our terminology for the “in *vitro”* and “*in vivo”* simulations: “in *vitro”* refers to “a single point injection, as per patch clamp experiments”, and “in vivo” refers to “multiple synaptic-like activity”. In this way we emphasize the significant difference whether the driving input was a single point or multiple synaptic sites.

## Results

### Dendritic morphology determines gain modulation

We begin with a modified ball-and-stick model, comprising of a single soma and approximately126 dendritic sections with only passive properties. We systematically rearranged the dendrites to vary polarity via the number of primary branches *n_p_* and the branching patterns via the number of sister branches at each bifurcation *n_b_* of the resulting arbor (see Methods) such that they were symmetrical around the soma (Fig. 1A). The total number of branches is controlled by the number of bifurcation stages (*n_ℓ_*, this number includes the creation of primary branches). This enabled us to generate 20 distinct dendritic morphologies. As the number of sections remained approximately constant, thus fixing the spatial extent of the dendritic arbor, this allowed us to identify if a neuron’s capacity for gain modulation could be determined by the dendritic arrangement, independently of total dendritic area, and if so, to establish which specific morphological features contribute. Effects of the total dendritic area have been investigated in [29].

**Fig 1.**
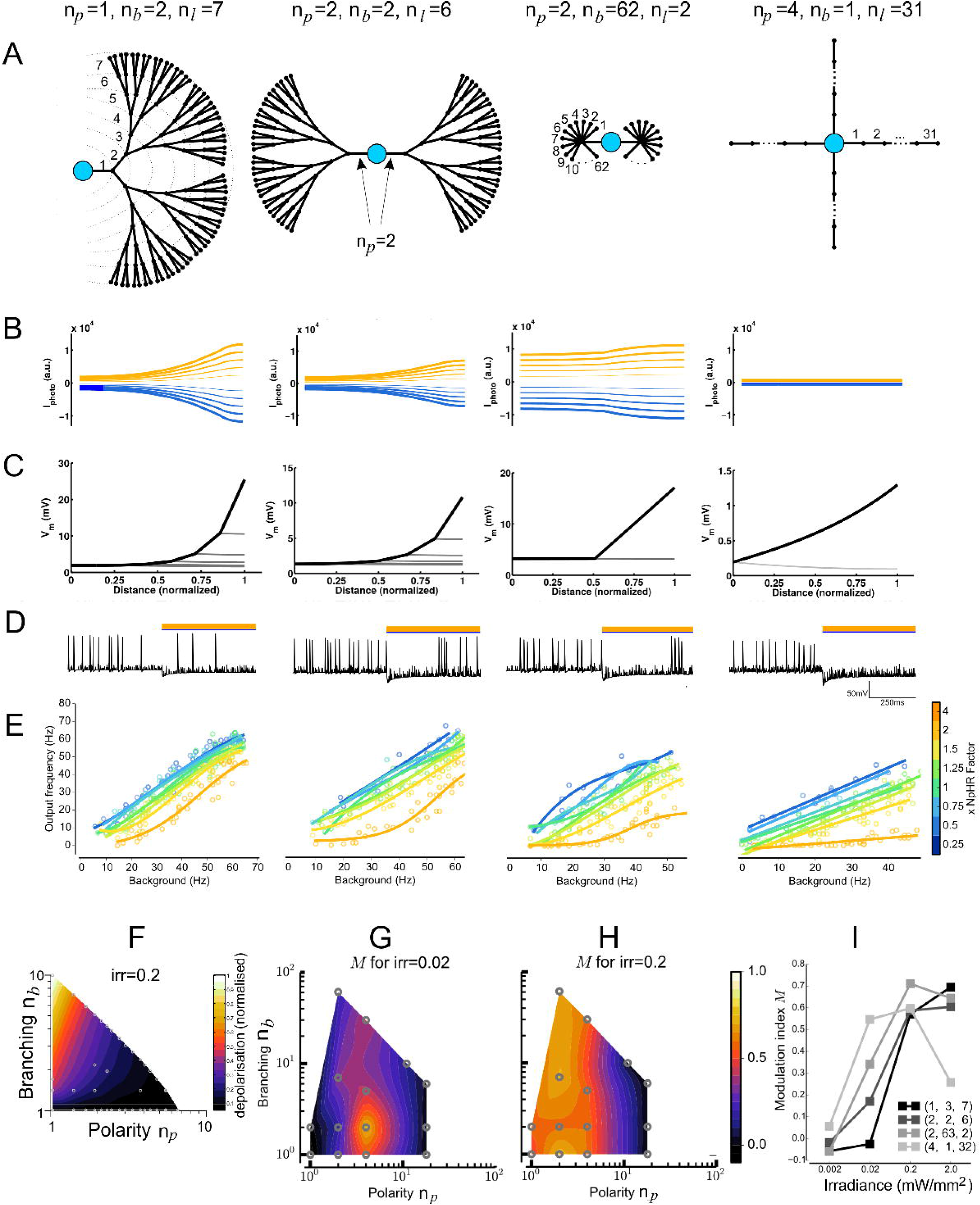
Gain modulation in model neurons with varying dendritic morphologies. (A) The four examples of dendritic morphologies that we examined here, which represent some typical morphological characteristics (left to right): (i) unipolar with moderate branching (n_p_ = 1, *n_b_* = 2, *n_ℓ_* = 7, total number of branches Ntotai = 127, see eq. (2)), (ii) bipolar with moderate branching (*n_p_* = 2, *n_b_* = 2, *n_ℓ_* = 6, *N*_total_ = 126), (iii) multipolar arbor with extensive branching (*n_b_* = 2, *n_b_* = 62, *n_ℓ_* = 2, *N*_total_ = 126) and multipolar with no branching (*n_p_* = 4, *n_b_* = 1, *n_ℓ_* = 31, *N*_total_ = 124); (B) The photocurrent measured at each compartment along the dendrite from soma to a terminating distal location, for four neurons, for inhibitory (orange) and excitatory (blue) photocurrents of different amplitudes; (C) Steady-state response for single injection of fixed amplitude at a distal location. Tracing the voltage for each dendrite section from injection site to soma (black), as well as sister branches and other primary branches (grey), shows the attenuation also varies with dendritic morphology, and is consistent with previous studies [30]; (D) Spiking responses (shown in panel F) were calculated by driving the neuron at a single distal dendritic section, and measuring the firing rates for unilluminated and illuminated cases - an example of the effect of using inhibitory photocurrents (x=4, see (E)); (E) Spiking frequency with photoactivation vs. background frequency (i.e. spiking rates prior to photoactivation) for the four example neurons showing no gain modulation, to increasing amounts of gain modulation, for *irr* = 0.02mW/mm^2^, for different ratios of NpHR and ChR2 illumination strengths from 1:4 to 4:1. No other conductances were changed.; (F) The net depolarization measured at the soma, for identical irradiance strength, across dendritic configurations. The x-axis: normalised values for Polarity (*n_p_*), y-axis: normalized values for Branching (*n_b_*).; (G) *M* values for a subset of dendritic configurations and *irr* = 0.02 mW/*mm*^2^, with *M* > 0 indicating gain modulation. Neurons with multiple primary dendrites and a moderate degree of branching displayed the most modulation. (H) Increasing the illumination strength by an order of magnitude (*irr* = 0.2mW/mm^2^) increases the region corresponding to morphologies that have the most modulation. (I) Modulation as irradiance is increased, for the four example neurons, shows a clear relation between matching irradiance strength to dendritic morphology in order to maximize gain modulation.

The contribution of branching has been investigated previously, and found to attenuate both voltage signals [30] and membrane resistance [20]. To understand the interaction between neuron-wide activation (as provided by the photocurrent) and a single point input (such as current injection or presynaptic input), we evaluated first the steady-state response of the abstract models when photocurrents were induced throughout the entire dendritic arbor, before considering separately the current injection in a single distal branch.

Excitatory and inhibitory opsins (ChR2 and NpHR, respectively) were included throughout the dendritic tree in addition to the soma, and generate excitatory and inhibitory photocurrents when photoactivated. We set the opsin expression (defined by gphoto in eq. (1)) to be inversely proportional to the area of each compartment, and fixed the irradiance to be equal across the entire neuronal surface, thus ensuring the resulting photocurrent induced for each section is constant. Measuring the net photocurrent locally along the length of the dendritic arbor while shifting the ratio between NpHR and ChR2 (*xNpHR*) whilst keeping all other conductances constant, different dendritic morphologies summate the photocurrents such that the voltage measured at the soma is influenced by branching and polarity (Fig. 1C), where morphologies with no branching show a small amplitude for the photocurrent. This relationship between net amplitude and branching was consistent across all arbor shapes we tested (Fig. 1B), providing the first indication that different neuron types will sum the photocurrents differently.

Like [30], we injected current at a single point on a distal, terminating branch and measured the membrane voltage across the path between the input site and soma, along with sites at sister branches, which indicated the amount of voltage attenuation that occurred without photocurrents included (Fig. 1D, note differing voltage scales). For a fixed amount of current injected on a single terminating branch, the perturbation when measured at the soma followed an identical trend as to that observed for the photocurrents in Fig. 1B, due to the symmetry of the dendritic configuration, varying however in magnitude. This suggests that the magnitude of depolarization from photoactivation has to be matched to that obtained from the point input; mismatch will result in the neuron’s output being determined by the dominant term. Thus, if there is a fixed point input, this requires the amount of photoillumination to be matched to the dendritic morphology. We additionally measured the depolarization when both photoillumination and current injection were included, and found that it was a linear sum of the responses we observed separately for both types of input, as expected as there were only passive ion channels in the dendrites.

Whether the input was dendrites-wide or a single point, these dual methods of driving the neuron illustrate their respective effects: that for both methods, depolarization is largest and most effective for branched structures. For sustained whole cell photoactivation, the induced photocurrent acts to raise or lower the effective resting membrane potential, upon which the depolarization from a single (or multiple) distal point can further drive the membrane at the soma to threshold. These results also indicate that the effect of the photoactivation can dominate the neuron’s response if not matched to the relative level of activation induced by current injection at a single point.

To contrast and compare these results with the impact of the dendritic tree when the input is located at the soma, we investigated the four characteristic dendritic morphologies from Fig.1A (Fig . This situation has been previously extensively investigated, e.g. Eyal et. al. [29], and was found that the shape and size of dendritic arbors strongly modulate the onset of action potentials by regulating the impedance load attached to the soma. The dendritic tree acts as a current sink in this case, and its impedance affects the soma depolarisation.

We then quantified the transient response by driving the neuron with spiking input, mimicking excitatory postsynaptic potential (EPSP) events included at set of locations at a terminating branch distal to the soma. By changing the rate of presynaptic events and then measuring the neuron’s firing rate we get the background firing rate when not illuminated, before repeating with irradiance (Fig. 1E), we were able to measure the gain of the neuron while varying the E:I balance by changing the ratio of ChR2 to NpHR (Fig. 1F). For *irr*=0.02 mW/mm^2^, we found that gain modulation was achieved in a subset of dendritic morphologies, marked by an increase in the gradient of the response as opsin activation moved from being dominated by inhibition to excitation (Fig. 1F).

To identify whether there was a consistent trend between dendritic configuration and gain modulation, we define a new measure we term the gain modulation index (*M*), as the relative change in gradient of the response curves (from Fig. 1F) when dominated by ChR2 and NpHR respectively, i.e. *M* = (*θ*_*ChR2*_ – *θ_NpHR_*)/*θ_balanced_* for the difference in angles for the two responses (the slope for *θ_balanced_* is approximately tan(*θ*_balanced_) ≈ 1). An *M* ≃0 indicates that no gain modulation occurred, whereas increasing *M* indicates an increasing degree of gain modulation. We found that there was a small region for which modulation was substantial, and correlated to dendritic structures that were multipolar with a small degree of branching (Fig. 1G, point *n_p_* = 4, *n_c_* = 2; note that a discrete set of measurements is additionally presented as a continuous colourmap for the purpose of better visualisation). Following our earlier indication that photoactivation has to be matched to dendritic structure, we measured the modulation for irradiance values an order of magnitude smaller and greater. When *irr*=0.002 mW/mm^2^ we observed no gain modulation for all dendritic configurations (not shown), as their responses were dominated by the current injection. Increasing the *irr*=0.2 mW/mm^2^ expanded the region of dendritic configurations which displayed gain modulation, which was now most prominent for bipolar morphologies (Fig. 1H). To obtain better intuition as to how irradiance affected modulation, we charted *M* for our four example neurons over four magnitudes and observed a clear trend, with preferred irradiance values for which a neuron will display maximal modulation (Fig. 1I). For *irr=2* mW/mm^2^ some configurations (e.g. (4,1,31)) start entering tetanic stimulation for full illumination with ChR2, for which the modulation M starts decreasing.

### Pyramidal cells are gain modulable

Following the predictions made by our abstract models, we investigated whether these principles still hold for detailed neuronal models, using a highly detailed Layer 5 pyramidal cell (Fig. 2A), previously published in [31]. Its reconstructed morphology is roughly bipolar with moderate branching, which, from the abstract models we tested, demonstrates a strong capacity for gain modulability. However, the model also contained 9 additional ion channel types heterogeneously distributed throughout the soma, apical and basal dendrites. These included multiple variants of Ca^2+^ and Ca^2+^-gated channels, which introduced **non-linearities** as well as significantly longer time constants, which may alter the capacity of a neuron to generate spikes and thus indirectly alter its capacity for gain modulability.

**Fig 2.**
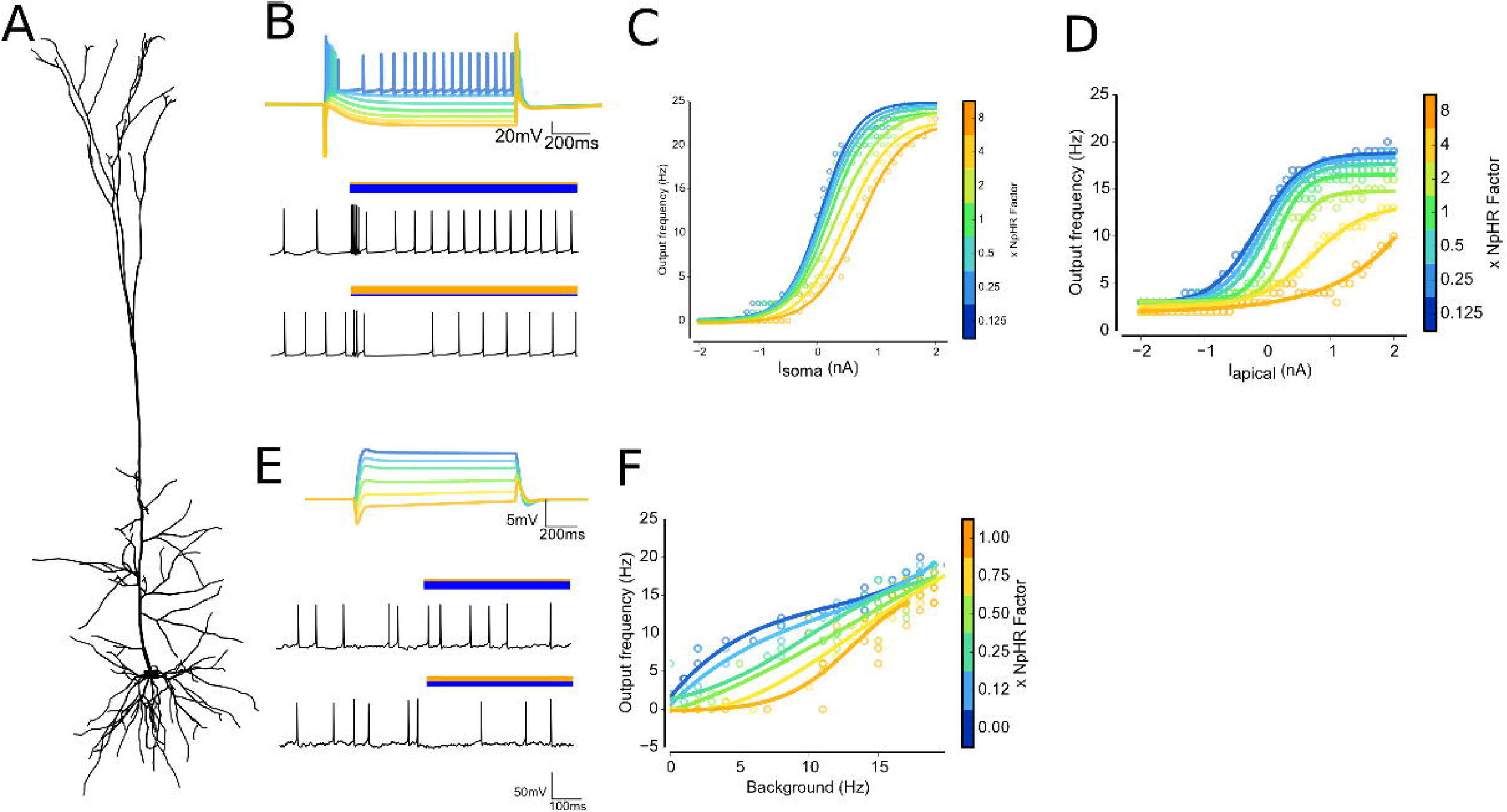
Simulated in *vitro* and *in vivo* responses for a pyramidal cell show gain modulation. (A) Layer 5 pyramidal cell, taken from [31]; (B) *(top)* Soma membrane voltage for the resulting net photocurrent as xNpHR is increased for irradiance=1mW/mm^2^ before current injection. *(bottom)* voltage traces following current injection while ChR2 and NpHR are photoactivated (bars indicate optical activation and relative strength of opsins); (C) Injecting current at the soma, mimicking *in vitro*-like input shows no gain modulation; (D) Moving the injection site to a location on the apical dendrite showed strong gain modulation; (E) *(top)* Same as panel (B, top) but for irradiance=0.02mW/mm^2^. *(Bottom)* Voltage traces after driving the neuron with multiple presynaptic EPSP and IPSP events, with photoactivation dominated by ChR2 and NpHR; (F) *In* vivo-like input showed moderate gain modulation, with saturation for larger input values. Background rates were calculated by varying the rates of presynaptic events and observing the period prior to photoactivation;

To reproduce experimental tests, we began by driving our L5PC by injecting current at the soma, in a similar manner to a typical *in vitro* electrophysiological experiment, and compared the firing rates upon illumination against the background firing rate (Fig. 2B). This revealed that IF curves were co-located (Fig. 2C), indicating no gain modulation. However, this was consistent with findings from our abstract models where we observed that gain modulation was site specific for the driving input. Consequently, we moved the injection site to a distal location on an apical dendrite. This time, we observed clear changes to the gradient of the IF curve as increasing amounts of current were used to drive the cell while varying the E:I balance (Fig. 2D).

While this demonstrated that the gain of this pyramidal neuron may be modulated in an *in vitro* scenario, neurons *in situ* are instead driven by thousands of excitatory and inhibitory synaptic inputs located throughout their dendritic arbor. Thus we repeated our simulation, but changed the input to mimic PSPs, by identifying 384 sites for excitatory inputs, and 96 sites for inhibitory inputs, throughout the apical and basal dendrites (Fig. 2E)-two bottom panels. We observed that gain modulation was still clearly evident (Fig. 2F), although the firing rates saturated for input firing rates greater than 20 Hz.

### Divisive modulation in stellate neurons - but only *in vitro*

To examine the effect of dendritic morphology, we also investigated gain modulation in stellate cells, which are also present within cortical circuits, but whose morphology is very different from pyramidal cells. We used a Layer II hippocampal stellate cell model previously published by [32], based on reconstructions from [33]. Morphologically, it is multipolar with a small degree of branching (Fig. 3A), which places it near to the abstract models for which we observed little to no gain modulation.

**Fig 3.**
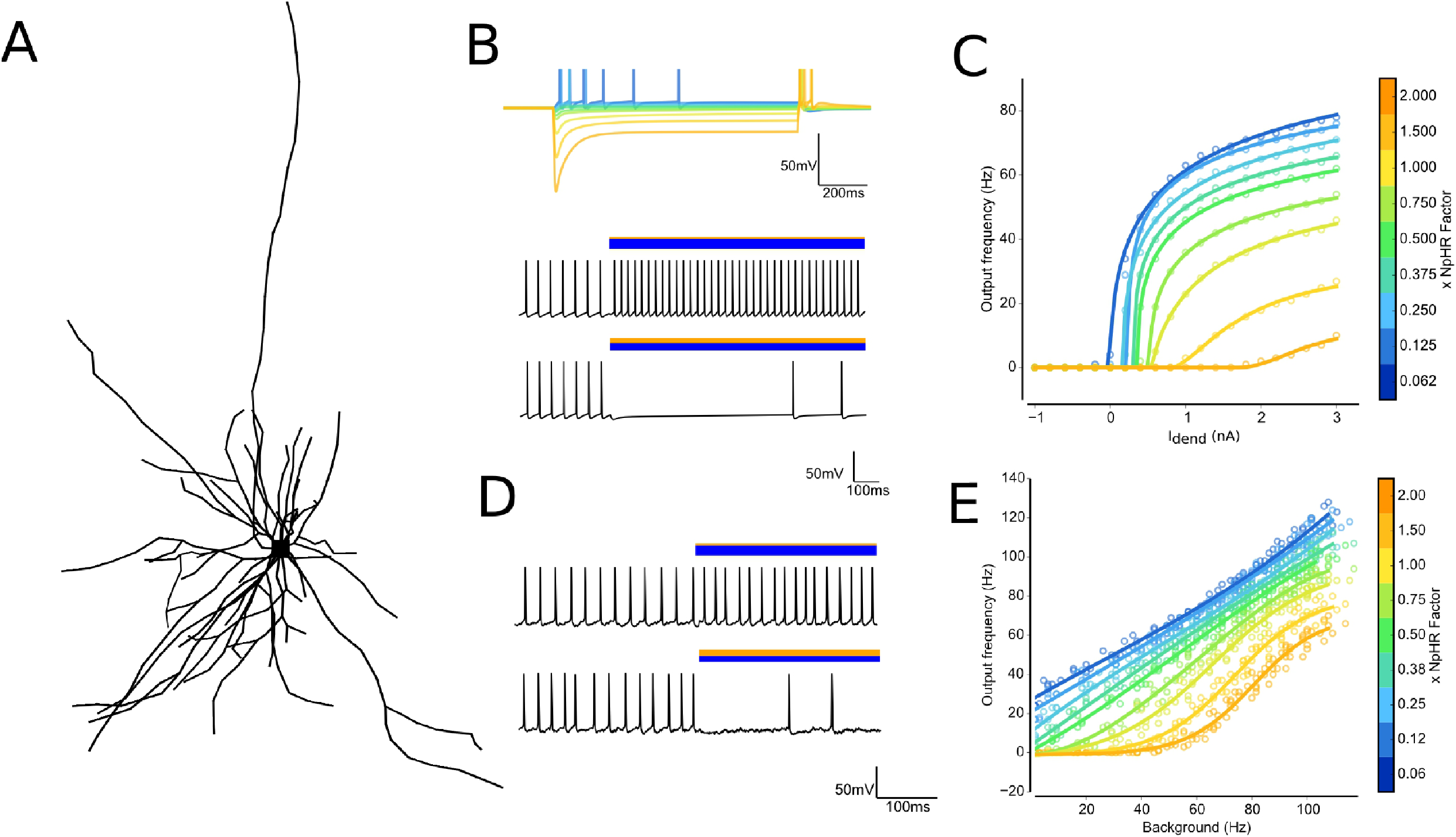
Gain modulation in stellate cells. (A) Layer 2 hippocampal stellate cell morphology, reproduced from model of reconstructed cell [32,33]; (B) *(top)* Membrane voltage for net photocurrent as xNpHR is varied for an irradiance of 0.07 mW/mm^2^. Current injection was fixed at a location on a dendrite more than 100*μm* from the soma. *(bottom)* Voltage traces, recorded at soma, for *I_dend_* = 1.2 nA respectively, following current injection while ChR2 and NpHR are photoactivated (bars indicate optical activation and relative strengths of opsin strengths); (C) IF curves for current injection indicates divisive gain modulation; (D) Voltage traces after driving the neuron with multiple presynaptic EPSP and IPSP events spread throughout the dendrites, with photoactivation dominated by ChR2 and NpHR respectively; (E) IF curves for *in vivo*-like input reveal no gain modulation (taken from multiple independent realizations for presynaptic spiketimes, *n*=5).

Unlike L5PCs, the response to an in *vitro* input of injecting current at a dendritic location (Fig. 3B) revealed that stellate cells do perform divisive gain modulation (Fig. 3C). In the case of current injection at the soma the same effect was observed as for the L5PC cell shown in Fig. 2C: an approximately co-linear response (Fig.).

We then drove the cell by supplying synaptic inputs throughout the dendritic tree to mimic *in vivo* conditions (Fig. 3D), and observed no gain modulation but rather a linear shift as *xNpHR* was varied (Fig. 3E). This suggests that while stellate cells presumably play an important role within the neural circuit, the gain of their input-output functions is unlikely to be modulated *in vivo*, but are instead likely to be subject to shifts in overall excitability through changes in the amount of excitation or inhibition.

### Shifting E:I balance produces smooth gain transition and preserves subthreshold dynamics

From our results, it is clear that for some neurons, such as pyramidal cells, it is possible to retune their output by applying whole-field photoactivation. However, by measuring the output firing rate, we ignore spike train structural characteristics such as the timing of the spikes. To examine how spike timing was affected by optogenetically altering the balance of excitation to inhibition, we considered a L5PC’s spike train in response to frozen noise input for an *in vivo*-like scenario and define a period during which we wish to increase or decrease the firing rate while the driving input remains fixed (Fig. 4A). High-level illumination, which is commonly used experimentally, dramatically reshapes the spike train as the membrane potential is either completely hyperpolarized or the neuron fires with a high-frequency, regular rate (Fig. 4B). Thus while this technically reprogrammes the gain of the cell, it does so by artificially rewriting the output spike times.

Instead, preserving subthreshold dynamics that arise from the hundreds of presynaptic events should allow the spiketrain characteristics to remain naturalistic, retaining rather than overwriting existing information processing functionality. To test this, we used a significantly lower level of illumination and found that the resulting spiketrains are qualitatively similar to the original response (Fig. 4C). To what extent can we perturb the neuron through external photoactivation before spiketrain characteristics are destroyed?

To quantify how the intrinsic spike timing of a neuron is altered by increasing levels of optogenetic activation, we measured the interspike interval (ISI) and then calculated the coefficient of variation of the interspike interval sequence (CVISI), and the Fano factor (FF), which describes the variance of the spiketrain normalized by the mean firing rate. We compared the CVISI and FF during the period where the firing rate was altered, as both the strength of illumination and the balance of excitation to inhibition was varied, for different levels of intrinsic activity. We observed that FF (Fig. 4D) and CVISI (Fig. 4E) could be maintained in the same range as the unperturbed spiketrain, but that this was dependent on level of illumination, suggesting that the artificial drive has to be matched to the level of the input the neuron already receives. The best matched level was for *irr* = 0.002mW/mm^2^, which closely matched the intrinsic CVISI and FFISI values of the neuron. Using this irradiance, we observed a smooth transition from the original response as the optogenetic drive moved from NpHR-dominated to ChR2-dominated (Fig. 4F).

**Fig 4.**
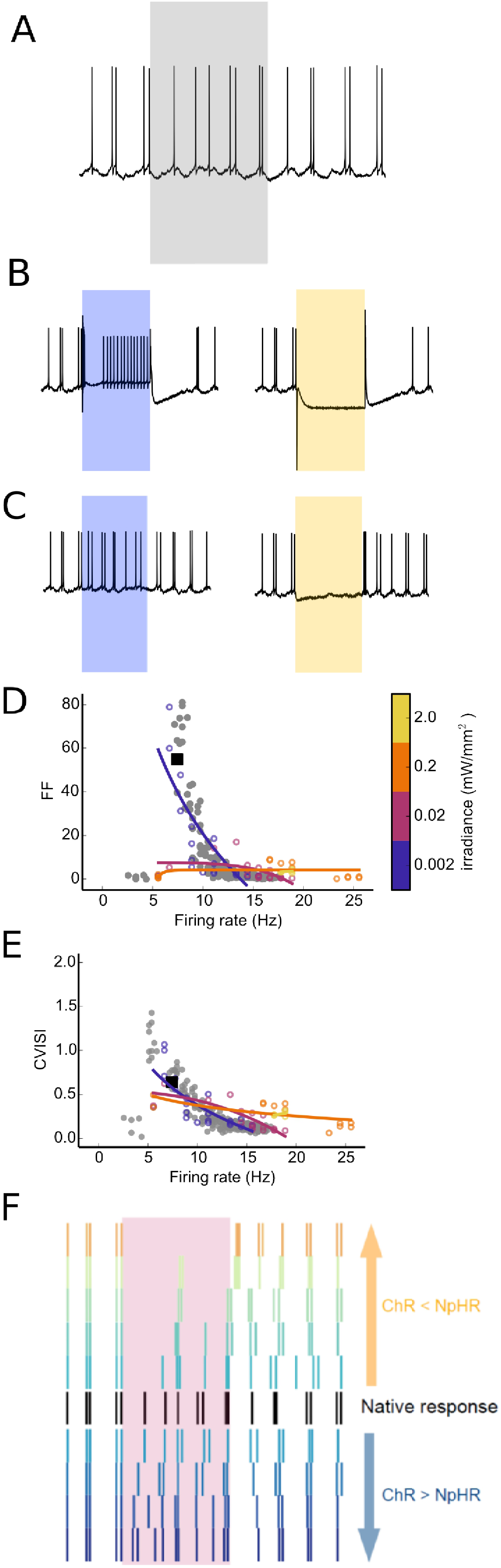
Subthreshold responses are preserved by low-level illumination. (A) The native response of L5PC when driven by presynaptic inputs of a fixed rate. Grey background indicates the time during which we wish to increase or decrease the firing rate; (B) Voltage traces V_soma_ when driven by high-level illumination (irr=2mW/mm^2^) for ChR2 (blue) or NpHR (orange) show clear increase or ablation of the firing rate. However, the membrane potential and spike train characteristics both during and post photoactivation are not naturalistic; (C) Instead, low-level illumination preserves subthreshold dynamics, minimally interfering with the subthreshold responses both during and post the activation periods (irr=0.002mW/mm^2^); (D,E) Quantifying spiketrain perturbances using the Fano factor (FF) and coefficient of variation of the interspike intervals (CVISI) during photoactivation as illumination strengths were increased. The FF and CVISI of the original response for firing rates as the presynaptic rates were varied (grey dots) as well as the sample plot shown in A (black dot) indicate the natural distribution of FF of the neuron’s response. Optical activation of the neuron when driven by presynaptic inputs over a range of xNpHR values while varying illumination strengths show that lower illumination strengths (blue) most closely match the FF of the natural cell and therefore minimally disturb the spike train; (F) The spike trains during the illumination period as xNpHR is varied for low-level illumination (irr=0.002mW/mm^2^) shows a smooth continuum of responses, as well as minimal disturbance to the post-stimulation spike trains.

### All dendritic subdomains are required for gain modulation

Our findings for both biophysically detailed and abstract models demonstrate that dendritic morphology greatly contributes to determining a neuron’s capacity for gain modulability. Up until this point, we have only considered scenarios with equal illumination for every dendritic subdomain. Experimentally, however, this is not guaranteed due to unequal expression of opsins throughout the cell membrane as well as uneven light scatter as photons move through tissue. Thus we investigated how gain modulation was affected when dendritic subdomains were unequally photoactivated. Mechanistically, this is relevant for gain modulation as synaptic input to a neuron is not likely to be uniformly distributed throughout the entire dendritic tree, but may instead be organized by presynaptic origin [21,34]. Could it be that such organization is present to allow the coordinated activation of dendritic subdomains, which is required for modifying the neuron’s output?

We began by examining partial illumination in abstract models, illuminating only one dendritic subdomain (pole) to examine how this altered gain as ChR2:NpHR was varied. We first want to pick the case with the strongest modulation gain during full illumination and we find that by looking at results in Fig. 1H): that is a bipolar model *n_p_*=2, *n_b_*=6, now illustrated in Fig. 5A) and the output frequency is shown in Fig. 5B). By illuminating only one pole instead (Fig. 5A), we observed that partial illumination abolished gain modulation (Fig. 5C). Measuring the voltage along both the illuminated and non-illuminated branches revealed that during partial illumination, the non-illuminated pole/branch acts as a current sink.

In this scenario, only 50% of branches were illuminated: perhaps gain modulation was still possible with an increased but incomplete set of dendritic subdomains? To test this, we need a multipolar abstract neuron (rather then the bipolar shown in Fig. 5A) and we chose one with *n_p_*=4 (and *n_b_*=2, *n*_l_=5), and successively activated additional subdomains (poles) until all branches were illuminated. We found that M increased as successive poles were illuminated (Fig. 5D). As this principle would hold for all dendritic morphologies, our findings demonstrate that partial illumination incapacitates gain modulation and illustrates that coordinated activation between dendritic branches is necessary for full gain modulation.

**Fig 5.**
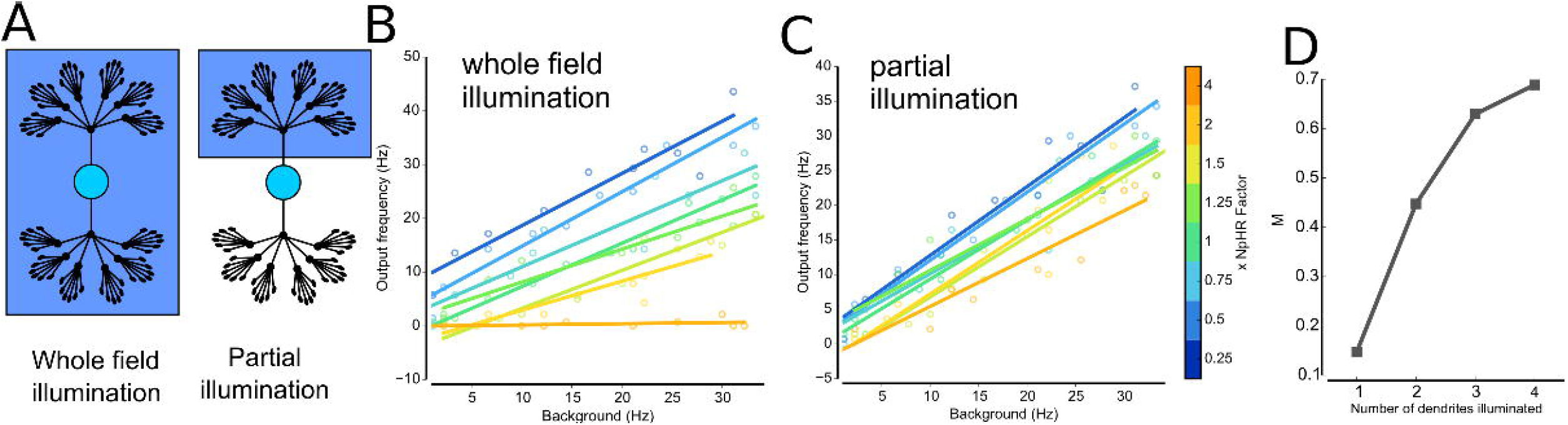
Partial and graded illumination in abstract neurons. (A) Full (left) and partial (right) illumination; (B,C) Response IF curve for a neuron with *n_p_*=2, *n_b_*=6, *n_ℓ_*=3 and irradiation = 0.2 mW/mm^2^ that has strong gain modulation when all branches (excluding the soma) are illuminated (B), but partial illumination nearly abolishes gain (C). (D) Measuring the modulation index M as successive illumination of poles reveals that gain modulation rapidly decreases as fewer branches are illuminated for a neuron with *n*_p_=4, *n_b_*=2, *n_ℓ_*=5 and irradiation=0.02 mW/mm^2^.

We then tested partial illumination in our detailed neuron model of a L5PC by targeting the apical dendrites, reflecting a realistic scenario in which light from a superficially located source would be more likely to penetrate the apical rather than basal dendrites (Fig. 6A). Similarly to abstract models with two primary branches, we found that partial illumination abolished gain modulation in L5PC when driven by current injection at a site in the apical dendrites (Fig. 6B).

**Fig 6.**
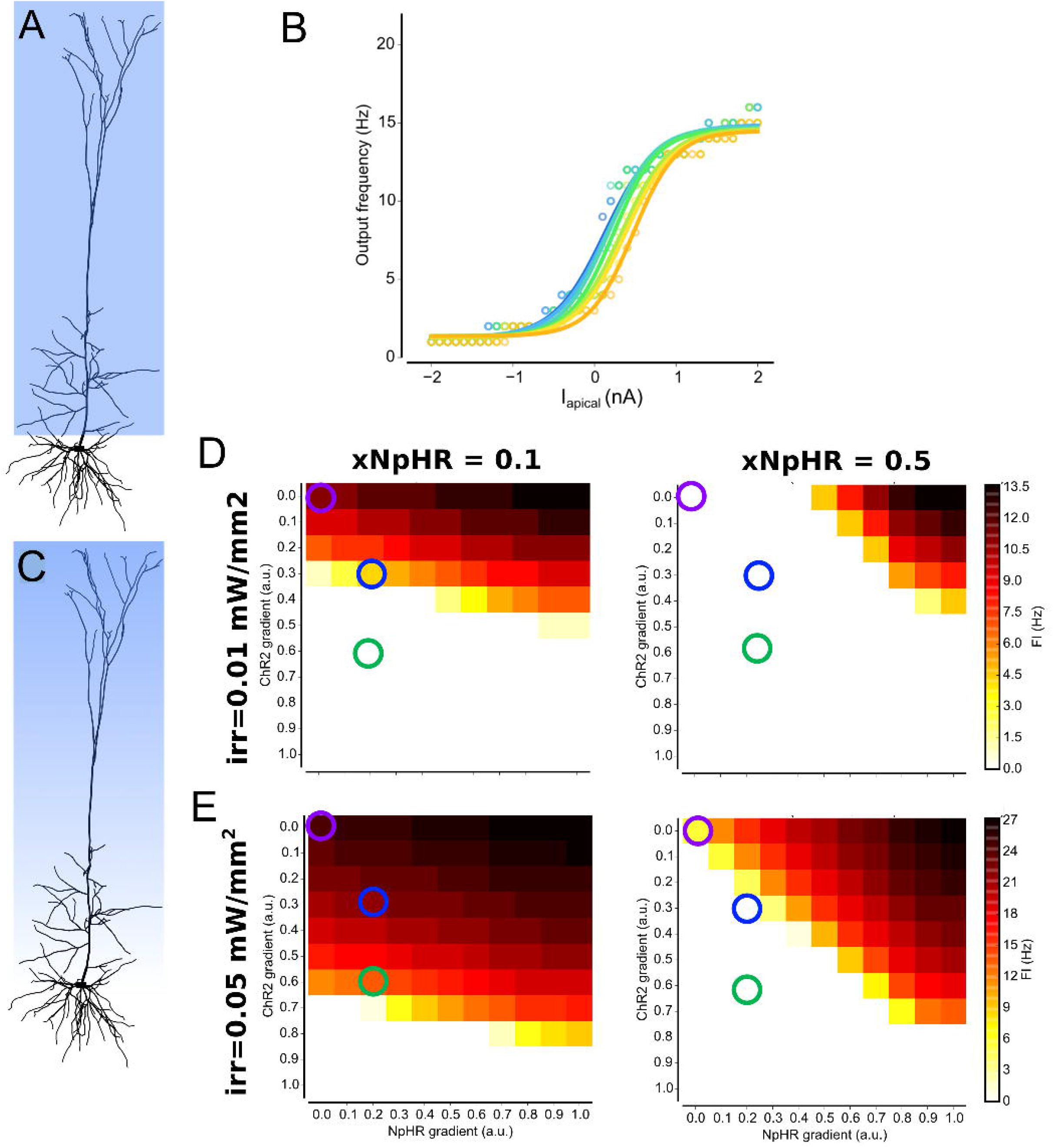
Partial and graded illumination in L5PC. (A) Partial illumination in L5PC model, where the apical but not the basal dendrites are illuminated; (B) The resulting IF curve for partial illumination shows no gain modulation for same input as used in Fig. 2D, where full illumination showed strong gain modulation; (C) Graded illumination over L5PC, from superficial layer at apical dendrites; (D,E) Graded illumination for fixed xNpHR and irradiance, for different combinations of graded illumination. For both axis zero represents no grading (i.e. uniform irradiation of complete apical tree), and one represents maximum grading which goes from the nominal irradiation at the top to zero irradiation at the bottom. The responses for no graded illumination (purple circles) are moderately reduced when ChR2 is scattered slightly more than NpHR (blue circles), and nearly abolished when ChR2 is scattered significantly more than NpHR (green circles).

A more realistic experimental scenario is one in which the likelihood of photons scattering rises with increasing depth, corresponding to a continuous gradient for the effective irradiance that decreases with distance from the surface (Fig. 6C). Furthermore, as opsin activation occurs by illumination using a wavelength that is normally chosen optimally for each opsin (subject to available laser lines), and longer wavelengths proportionally penetrate distances, we examined the penetration gradients for ChR2 and NpHR independently (Fig. 6D). Previously, we had only considered full-illumination of both ChR2 and NpHR with no graded illumination, which was equivalent to a gradient value of 0.0 (Fig. 6D, top-left corner, purple circles). Now, by fixing *xNpHR* and irradiance while activating the L5PC without any additional driving stimulus, we could chart how independently varying each gradient impacted on the firing rate.

We observed three trends that are consistent with our earlier observations: (i) increasing the *xNpHR* factor increased the contribution of the NpHR gradient, which decreased the firing rate; (ii) a ChR2 gradient=0.0 (signifying full ChR2 illumination) and a NpHR gradient of 1.0 resulted the largest firing rates; (iii) higher irradiance values led to higher firing rates, increasing the range of ChR2 gradients for which the neuron fired. However, we were interested in cases that correspond to the realistic scenario in which longer wavelengths penetrate through tissue further. As the preferential activation wavelengths for ChR2 and NpHR are λ = 475nm and 590nm respectively, this manifests as a bias towards NpHR-dominated regimes. Introducing a small degree of graded illumination reduced the firing rate; the neuron was further silenced by increasing the NpHR gradient from a slight bias (Fig. 6E, blue circles) to a significant relative difference between ChR2 and NpHR gradients (Fig. 6E, green circles).

The modulation by graded illumination was ubiquitous, although dependent on the irradiance, *xNpHR* value and scatter gradients for each wavelength. Experimentally, these effects can be easily overcome by prior calibration to compensate for the effects of scattering, but serve to highlight the sensitivity of a neuron to deviations from unequal innervation.

## Discussion

### An ideal dendritic morphology for gain modulation?

Previous work [12–14,16] examined what input properties are required to alter the output gain of the neuron. Critically, these studies took a somatocentric viewpoint, concentrating on the output of the neuron for a given input, but bypassing the computation performed by the neuron itself. In this work, we addressed the contribution of the dendrites directly, by considering how their configuration may help or hinder modulation of the neuron’s activity and thus explain why some classes of neurons, but not others, contribute to setting the gain in a neural circuit. We established that the configuration of dendrites can affect a neuron’s capacity for gain modulability, with a centrally placed soma and a moderate amount of branching being most receptive to gain modulation. As this shape closely matched pyramidal cells, this reinforces that their role within neural circuits is to act not only as the primary excitatory neuron but also as a key element in the setting of the gain of the circuit. Thus, in addition to the influence of dendrites on firing patterns [24,35] and their role in dendritic computation [36,37], our results demonstrate a new aspect to dendrites that directly relates the morphological properties of an individual neuron to its functional role within a network.

We explored the relation between a neuron’s dendritic morphology and capacity to alter its firing rate by using excitatory and inhibitory photocurrents locally input to each dendritic section. The use of photocurrents, as well as making the study relate more closely to putative optogenetic validation experiments, was intended to mimic the local excitatory and inhibitory currents induced by the numerous presynaptic inputs located throughout the entire dendritic tree, with the notable difference in that while postsynaptic potentials are transient, the photocurrents we induce were typically close to steady-state. Further input was additionally applied that mimicked *in vitro* or *in vivo* input. We made no specific assumptions as to the specific type of stimulus representation of the input, such as visual contrast [38], orientation [10] or other stimulus traits; our results hold for the general case in which a driving input at discrete set of location is modulated by neuron-wide distributed drive.

Using this framework allowed us to identify that dendritic branching and the relative location of the soma were the most important morphological characteristics, as dendritic branching allowed balance between saturation and attenuation while a centrally located soma avoided it acting as a current sink. As previously established in [30], these effects become crucial when considering the compounded local input that is applied to each dendritic section (here, the non-driving input i.e., the photocurrents). For L5PC neurons, the stratified output for illumination suggests that if net drive to the dendrite is able to sufficiently cover the entire arbor, then L5PCs will be gain-modulable independent of the location of the driving input, which has been shown to govern the input-output relationship for single inputs [18,39].

In this respect, our findings suggest that gain modulation should be achievable regardless of the specific input location. However, there is one critical caveat: that the input must be dendritic. The absolute abolition of gain modulation when the driving input was located at the soma in a pyramidal cell reinforces the role of dendrites in processing input and their contribution to modulating gain. It also highlights the difficulties associated with experimentally unraveling neuronal mechanisms which involve dendritic processing. While recording at the soma gives us an exact measure of the cells output, injecting input directly to the soma bypasses the dendrites, rendering their contribution invisible. Instead, techniques such as dendritic patching or extracellular drive are more suitable for this purpose, despite their respective technical challenge or lack of control for the number and locations of synaptic sites. The future development of holographic methods in combination with optogenetics provides potential solution to both limitations, although it is currently limited by the tradeoff between number of distinct sites that can be targeted and the frequency of their stimulation [40].

We note here that we attempted to use only passive conductances in biophysical models, in order to have better comparison with abstract neurons in interpreting the role of dendritic morphology, since their models include some active conductance (although in relatively low concentrations). However, due to their configuration and complexity we were unable to obtain normal neuronal responses i.e resting potential did not equilibrate in the experimentally observed range, due to different parameters required for the L5PC and stellate cells to operate in the passive regime.

### The importance of presynaptic targeting for full gain modulation

Quantifying the effect of partial illumination revealed that gain modulation also requires the participation of all dendritic subdomains. Removing background input from dendritic subdomains resulted in the unactivated arbors becoming a current sink, and reduced the ability of the neuron to modulate gain.

Tracing studies have hinted that within pyramidal cells in sensory areas, there is synaptic targetting with feedforward presynaptic input from the thalamus tends to synapse onto basal dendrites, while input from higher cortical areas instead connects within apical dendrites [34,41]. This arrangement of separate innervation to distinct dendritic subdomains would very easily allow for the same mechanistic process as we have observed here, whereby both feedforward and feedback connections are required for full gain modulation. Removal of one of these sources, such as the feedback input from higher cortical areas, would quickly act to shunt any background drive to corresponding dendritic subdomain, thus providing a mechanism for rapid switching between full gain modulation and no gain. As we observed that the modulation of gain approximately scales with the fraction of the dendritic subdomains that receive background driving input, the change between full gain and no gain can be most effectively controlled with two dendritic subdomains, as increasing the number of subdomains requires greater coordination between distinct input areas.

### Application and improvement of optogenetic protocols

An important feature of computational models of neuronal information processing is that they be experimentally testable. Traditionally, for biophysically detailed, compartmental models of neurons, this has involved making predictions that can be confirmed by intracellular recording (whole cell patch clamp or sharp microelectrode) experiments. However, recent years have seen new optogenetic experimental paradigms come to the forefront of neuroscience, which are likely to form the basis of many experimental designs to test principles of neuronal function. Computational modeling of neuronal function should incorporate simulation of experimental predictions made by the model; whereas in the past this was largely electrophysiological, this now includes both electrophysiological and optogenetic predictions. We envisage that computational optogenetic modeling is likely to assist in bridging the gap between computational and experimental studies in areas ranging from neuroscience [42, 43] to cardiac electrophysiology [44]. For this reason, in the current study we incorporated kinetic models of opsin into the biophysically detailed neuron models described here.

Optogenetic illumination protocols in current use can generally be classified as “hard control”, in which the output of a cell is written directly by using high levels of illumination to induce either spiking or hyperpolarization [45,46]. The problem with such approaches is that they effectively reprogram the output of the neuron, disrupting/eliminating the information processing operation that it is performing on its inputs. We suggest that a more refined method of optogenetic modulation would preserve the cell’s ability for its outputs to be affected by its inputs, but altering the gain of this input/output transformation. Our findings demonstrate the feasibility and support the development of such optogenetic control of individual neuronal gain. In this approach, using whole-field, low-level illumination allows for subthreshold dynamics to dominate, and the neuron remains driven by its presynaptic input, with the gain of its input-output function modulated by activation of a mixture of opsins. In the current work, we demonstrated that a combination of channelrhodopsin and halorhodopsin can provide a suitable opsin mix, with effect dependent upon target cell morphology.

For the general purpose of optogenetic gain control, step-function opsins (SFO) [47] and stabilized step-function opsins (SSFO) [48] may be suitable, as they do not required continuous illumination to be active, and have their suitability for loose control has already been demonstrated when driven by inputs located at the soma [47]. SFOs and SSFOs have already been proposed for use in the study of plasticity and homeostatic mechanisms during development. Our results support their suitability for application to gain modulation *in vivo* but also predict a restriction to their usage that will be dependent on the class of neurons to be targeted. More generally, our findings suggest that smaller, rather than larger, photocurrent amplitudes are desirable for the purpose of modulating gain.

Unequal or incomplete optical activation of the entire dendritic arbor also has significant implications for experiments that include optogenetics. We used optogenetics specifically as opsins are expressed on the surface membrane, and therefore can generate photocurrents locally within the dendrites. Experimentally, however, opsins may be non-uniformly distributed throughout the neuronal membrane, while optical point sources incompletely illuminate the entire membrane surface area, which is further compounded by scatter effects as light moves through tissue. We quantified the impact of optical scattering by examining how graded illumination alters the gain modulation curves of a L5PC, for the scenario when the scattering was equal but also for the more realistic scenario where it is unequal. For instance, ChR2 is activated at λ=475nm, while NpHR is preferentially activated at λ=590nm, which penetrates further through tissue. From our results, approximately equal attenuation for both wavelengths only slightly decreases the firing rate; as this imbalance increases and shifts towards longer wavelengths, we found a substantial decrease in firing rate due to this physical constraint that biases in favor of NpHR. Additionally, although ChR2 and NpHR have different peak absorption wavelengths, they both are activated over a larger range of wavelengths, and consequently there is low-level activation of NpHR at λ=475nm, further biasing dual activation towards NpHR-dominated regimes. These effects can be experimentally compensated for by calibrating curves as the ratio of ChR2 to NpHR is varied, but require explicit measurement for individual experiments.

The issue of precise co-activation of both excitatory and inhibitory opsins can be addressed by development of better dual opsins, as well as of better techniques for controled 3D illumination. Several options already exist, such as holographic spatially patterned illumination technology [49] and individually addressable LED micro-arrays [50].

More generally, identifying the impact of experimental effects is critical for the improvement and refinement of optogenetics. While optogenetics offers new possibilities for precise spatial and temporal targeting of distinct neural populations, practical hurdles such as optimally designing illumination protocols are more difficult to identify through experimental means. Additionally, as new opsin variants with differing kinetics becoming available, the task of identifying which opsin is best suited to match the intrinsic dynamics of a target neuron class becomes increasingly impractical to test. For these aims, the use of computational models of opsins will become increasingly significant [51], from the level of channel kinetics, to the level of a single neuron [52,53], and beyond to the level of the network.

## Methods

All simulations were performed in NEURON and Python [54]. The TREES toolbox [55] was used for steady-state analysis of injected current in abstract models, and NeuroTools toolbox [56] was used for spiketrain analysis. Our simulations were calibrated with experimental results for which we performed a rigorous fitting process to the experimental data (ours for ChR2 and ref. [57] for NpHR). Details of the processes can be seen on our Cloud Computing Portal – named Prometheus – which hosts all our computational tools for optogenetics (called PyRhO): http://try.projectpyrho.org/. This is a Python-based Jupyter Notebook that works in a web-browser. Within the user interface, functions for fitting parameters allow experimental data to be loaded from which parameters are extracted for 3, 4 and 6 state models. The tool offers nine different fitting algorithms to chose from, as well as post-fit optimization.

### Modelling co-activated opsins for dual control

Our model of a co-activated opsin utilizes our previously published 6-state models of channelrhodopsin-2 (ChR2) [43,52] and halorhodopsin (NpHR) [53]. A 6-state model was chosen for both ChR2 and NpHR, which includes two open states, two closed states and two inactivated states that are coupled together with by rate constants. Only the open state contributes to the generation of the photocurrent *i*_photo_ for each opsin type, which is calculated per compartment and is additionally proportional to the area A and maximal conductance for each opsin 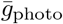 in combination with two terms related to the irradiance *ϕ* and the membrane voltage *V_m_*:

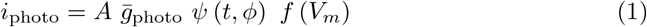

Critically, these models accurately capture the ion concentration kinetics and so allow accurate modelling of subthreshold dynamics, and can be tuned to provide a faithful reproduction of the temporal courses induced by opsin activation.

Throughout this work we refer to the ratio xNpHR as the relative strength of NpHR illumination in reference to a fixed value for ChR2 illumination.

### Abstract neuron models

Each of the abstract neuron models we created included a soma and approximately 125 dendritic sections that were arranged with varying degrees of branching. Dendritic morphology was described as the number of primary branches *n_p_*, the number of sister branches *n_b_* and the number of branching stages *n_ℓ_*. The total number of branches (*N*total) is given by:

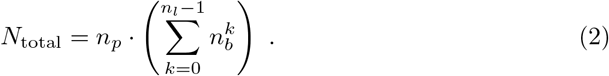

Altogether, we generated 31 different dendritic configurations that, despite their geometrical configuration, had approximately equal surface areas and volumes. Dendritic tapering was excluded to conserve total surface area.

For the biophysical properties, the values for the soma were *C* = 1pF, diameter=10*μm* and length *L*=10*μm*, while each dendritic section had parameters diameter=0.4*μm*, *C* = 2pF and length *L* = 50*μm*. All sections and soma had passive membrane properties (g_pass=0.00005, e_pass=-75mV) while the soma additionally had Hodgkin-Huxley channels, the NEURON built-in hh mechanisms were used, with conductances: gnabar_hh = 0.25, gkbar_hh = 0.1, gl_hh = 0.000166, el_hh = -60mV, ek = -70mV.

Each soma and dendritic section had ChR2 and NpHR models inserted, with constant expression throughout the dendrites and soma.

Constant driving input for the steady state response was modeled by injecting constant current in the last segment of a single distal segment, and normalizing the distance from total length to soma. Synaptic locations were chosen randomly from all distal sections. Synapses themselves were modeled using NEURON’s `ExpSyn` model, with input spiketimes drawn from independent Poisson process. As different dendritic morphologies had different electrotonic distances from synapse location to soma, synaptic weights were chosen where possible such that the output firing rate was approximately equal to the input firing rate, enforcing a loose version of synaptic democracy. For some arbor configurations, the length from soma to distal dendrites was greater than the electrotonic distance and a transient response could not be obtained. The dendrite tapering was not included in our simulation for abstract neurons, but it is included for two realistic biophysical neuron models. Inclusion of tapering for abstract neurons creates a problem of keeping approximately the same total dendritic tree area for various morphologies, which was the more relevant factor with respect to our results.

### Biophysical models

#### Layer 5 Pyramidal Cell

We used a previously published model of L5PC [31], ModelDB: no.139653 (cell: L5PCbiophys3).Both excitatory and inhibitory PSPs were distributed throughout the apical and basal dendritic trees. 160 synaptic sites were chosen (80 on the apical tree, 80 on the basal dendrites) at a minimum distance of 100 microns from the soma.

#### Layer 2 Hippocampal Stellate Cell

We used a model previously published by [32], based on reconstructions from [33], ModelDB: no.150239 (cell: stellate-garden). For in vivo input, synaptic locations chosen at random across the dendritic branches so that there were multiple sites that were equidistant. In total, there were 40 sites, at a distance approximately 100 microns from the soma.

*The total surface area* is approximately: i) for L5PC: the dendritic tree is approximately 60,000 *μ*m^2^ and the soma area is 1,131 *μ*m^2^, ii) Stellate cell: 30,000 *μ*m^2^ dendrites. For comparison, the corresponding values for abstract neurons are: 8,000 *μ*m^2^ for the dendrites and 314 *μ*m^2^ for soma,

#### Input to biophysical models

For both cells, ChR2 and NpHR models were inserted in the soma and dendritic subdomains as required. Additional input was provided either as a current injection at a single location, or as a barrage of inputs that created postsynaptic potentials, using neuron’s `ExpSyn` model. The spike times of each input site were generated as an independent Poisson process. The amplitude for excitatory PSP input was 0.5mV and inhibitory PSP 0.1mV for the L5PC, and 2mV and 6mV respectively for stellate cells. All synapses had a time constant of *τ*=0.1ms.

## Supporting Information

**S1 Fig.**
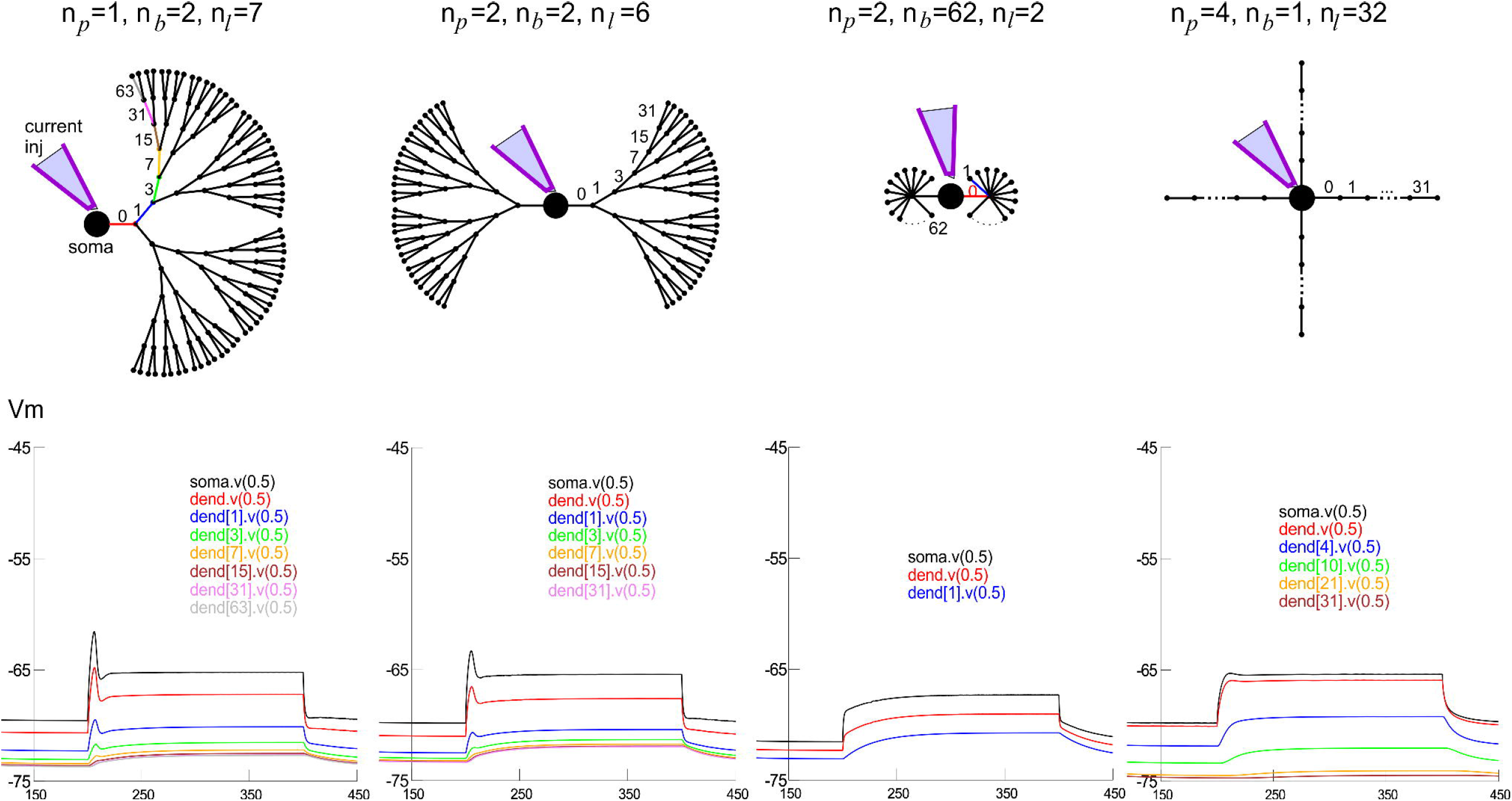
Injecting current at the soma in abstract cells shows membrane voltage changes dependent on the dendritic tree input impedance from the some side. (top) The four characteristic dendritic morphologies examined in Fig.1 and the positions of the dendrites that were chosen to represent different distances from soma to the endings of the dendritic tree. (bottom) The membrane voltage at the chose locations. For approximately the same number and surface size of dendritic arbours, the morphologies with lower input impedance will sink more of the injected current and hence cause less depolarisation at soma.

**S2 Fig.**
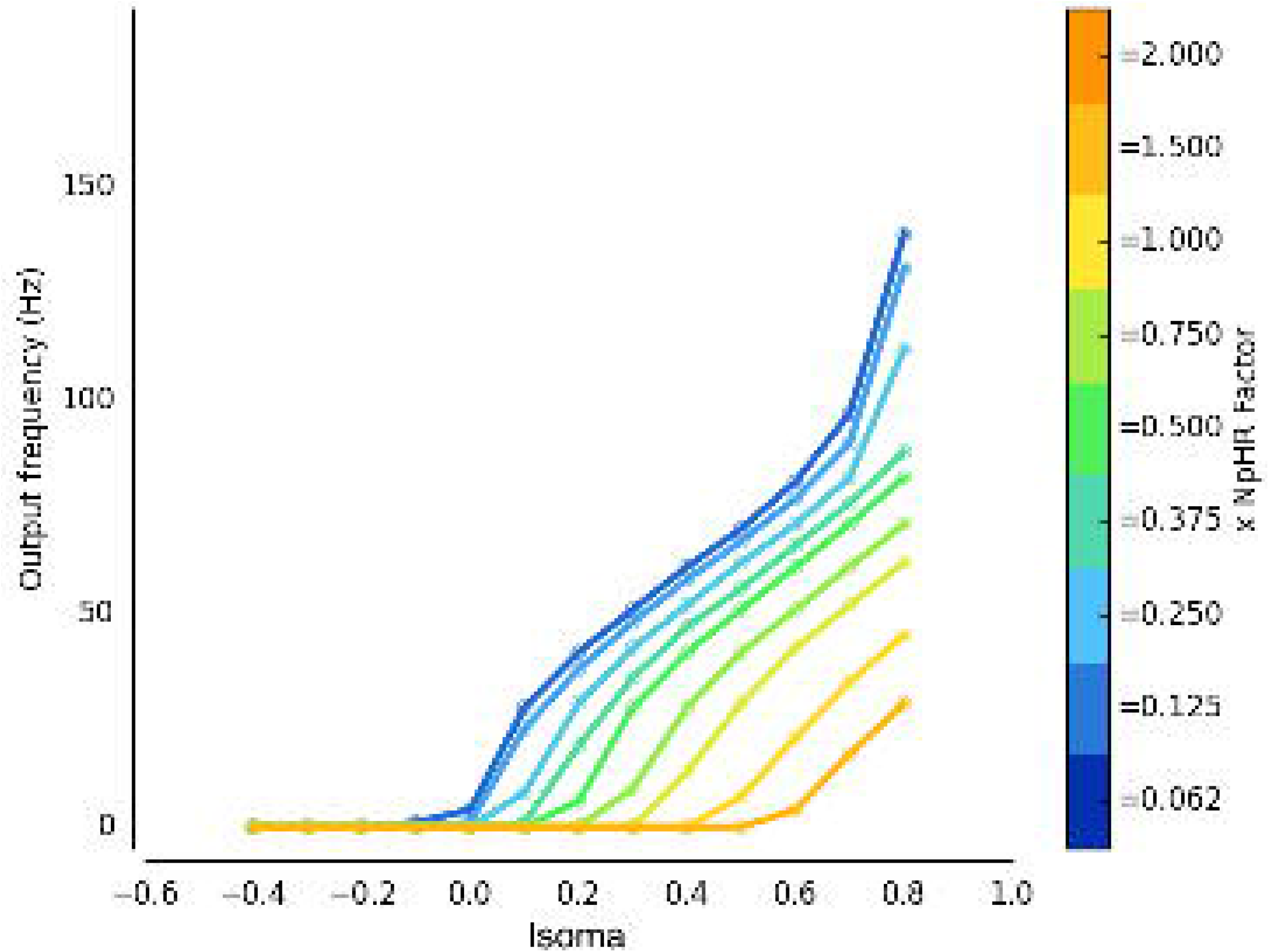
Injecting current at the soma in stellate cell shows no gain modulation. The membrane voltage for net photocurrent as *xNpHR* is varied for an irradiance of 0.07 mW/mm^2^ as it was in Fig. 3B, but with current injection occurring at the soma instead of dendrite. As observed for current injection at soma in pyramidal cell model (Fig. 2C), no gain modulation occurs.

## Acknowledgments

The authors thank Claudia Clopath for useful feedback.

